# Matrix-modulated Air-liquid Interface Microenvironments Reveal Architectural Control of Lung Tumor Phenotypes

**DOI:** 10.64898/2026.10.10.758195

**Authors:** Kardelen Yangin, Ece Öztürk

## Abstract

Lung adenocarcinoma evolves within a highly structured microenvironment shaped by air exposure, basement-membrane ligands and three-dimensional (3D) organization, yet standard in vitro models capture only fragments of this architecture. Here, we systematically disentangle how air-liquid interface (ALI), extracellular matrix (ECM) cues and dimensionality interact to regulate tumor epithelial states. Using five architectural configurations, we demonstrate that these parameters act synergistically to promote distinct morphological, transcriptional and inflammatory phenotypes not observed under conventional 2D or submerged 3D cultures. ALI broadly enhanced epithelial cohesion, progenitor activation, partial EMT programs and cytokine expression, whereas ECM embedding further induced ALI-mediated mesenchymal activation, matrix remodeling and invasive cytoskeletal organization. Notably, 3D ALI uniquely produced a hybrid alveolar-mucinous phenotype characterized by concurrent SP-C and MUC5B induction, alongside integrated epithelial, basal and mucosecretory identities. These emergent states reflect tumor-relevant plasticity and illuminate architectural controls over lineage infidelity, invasion and inflammatory activation. Together, these findings establish ALI-ECM-dimensionality intersections as critical regulators of lung tumor phenotype and provides a systematic architectural framework for designing more physiologically relevant in vitro lung cancer models.

## 1. Introduction

Lung cancer remains the leading cause of cancer-related mortality worldwide [1]. Non-small cell lung cancer (NSCLC) accounts for the majority of cases and adenocarcinoma constitutes the most frequent histologic subtype [2]. Adenocarcinoma commonly originates in the peripheral lung where the apical side of the epithelium is directly exposed to air while the basal side faces interstitial fluids and extracellular matrix (ECM), a quintessential air-liquid-interface (ALI) context which conventional submerged cultures fail to recapitulate [3]. Thus, honoring the microenvironmental conditions that the tumor unfolds with models representing epithelial polarity, the ECM and gas exposure is required for a better understanding of tumor cell behavior.

The airway epithelium is a specialized, multicellular barrier comprising basal progenitors, goblet and multi-ciliated cells which together maintain mucociliary clearance, host defense and tissue homeostasis [4]. Differentiation and function of these cells are profoundly affected by epithelial polarization and air exposure [5]. Accordingly, ALI culture on permeable supports has become the benchmark method for creating a pseudostratified, mucociliary epithelium *in vitro* with tight junction formation and transepithelial electrical resistance (TEER) [6,7]. Single-cell and organoid studies demonstrate that ALI remodels cellular composition and transcriptional programs toward ciliated lineages which underscores air exposure as an active driver of epithelial fate [8–10]. In parallel, mucins such as MUC5B and MUC5AC are canonical readouts of ALI-induced secretory differentiation which establish the polymeric framework of airway mucus with well-defined roles in both airway physiology and disease [11,12]. Despite this physiological backdrop, the application of ALI paradigms to *in vitro* lung cancer research has lagged behind, with most NSCLC studies conducted in submerged two-dimensional (2D) monolayers or three-dimensional (3D) matrix-embedded spheroids [13–15]. These models have been instrumental for mechanistic studies and therapeutic screening, however, they do not capture apical air exposure and polarity inherent to the respiratory epithelium.

Seminal work has shown engineered respiratory models which integrate ALI within microphysiological systems. “Breathing lung-on-a-chip” devices emulate the alveolar-capillary unit by combining microfluidic perfusion, cyclic stretch and ALI, enabling studies of nanoparticle absorption, barrier integrity and host responses under such dynamic conditions [16,17]. Incorporation of lung cancer cells onto pulmonary epithelium grown on such devices revealed key roles of biomimetic cues such as stretch application and showed its involvement in cancer cell growth as well as response to EGFR inhibitors [18]. Multilayered ALI cultures of NSCLC cells demonstrated increased chemoresistance compared to 2D monolayers and allowed aerosolized drug delivery using clinical nebulizers [19]. These models have also been adapted for fibroblast co-cultures revealing roles for cancer-stroma crosstalk in multidrug resistance [20]. At primary tissue level, ALI patient-derived organoids were shown to preserve endogenous immune infiltrates, stabilize tumor-immune architecture lost in submerged Matrigel cultures and allow modeling checkpoint inhibitor responses as well as capturing immune pharmacodynamics [21].

Along the ECM axis, the lung tumor microenvironment (TME) is defined by a basement membrane (BM) and interstitial scaffold enriched in laminin isoforms, collagen IV, collagen I/III, nidogens and heparan-sulfate proteoglycans, presenting distinct biochemical ligands that regulate epithelial polarity, adhesion, receptor activation and lineage specification [22]. These biochemical cues critically shape lung adenocarcinoma behavior via engaging integrins (i.e. a3b1, a6b4), discoidin domain receptors (DDR1/DDR2) and receptor tyrosine kinases through proteoglycan interactions, which together modulate proliferation, epithelial-mesenchymal transition (EMT), migration and plasticity [23–25]. Altered ECM composition in NSCLC has been shown to promote invasive and therapy-resistant phenotypes via signaling routes including COL10A1-DDR2-FAK, EGFR and TGFR [26–28]. ECM-dependent mechanical features can further amplify biochemical signaling, reinforcing pathways such as b1-integrin-FAK, EPHA2/LYN/TWIST1 and YAP/TAZ to induce EMT and invasion [29–31]. These studies underscore that the ECM is not a passive scaffold but an actively instructive biochemical and biophysical regulator of lung tumor phenotype, rendering the presence of BM-derived cues critical for engineered *in vitro* models.

Taken together, ALI and ECM represent two orthogonal axes that are essential for predictive modeling with distinct effects on epithelial organization and tumor phenotype, yet existing studies typically vary only one at a time. Although the shift from 2D monolayers to 3D cultures has improved physiological relevance, most 3D systems remain fully submerged, leaving the role of air exposure unexplored. A factorial assessment of ALI, ECM and dimensionality in regulation of lung tumor phenotype has not been explored in the field. In this work, we address this gap by systematically probing how air exposure, ECM cues and culture configuration intersect to re-pattern tumor epithelial states. Our findings reveal that these parameters impose distinct shifts in tumor epithelial organization yielding markedly different outcomes in growth, morphology, barrier function, lineage engagement, EMT and matrix remodeling. Our work demonstrates that lung adenocarcinoma cells adopt fundamentally different phenotypic regimes depending on the structural and biochemical cues presented in their microenvironments and offers design perspectives for building physiologically anchored in vitro respiratory tumor models and enabling more faithful evaluation of therapeutic interventions.

## 2. Results

### 2.1 Distinct tumor epithelial states emerge from modulation of air exposure, ECM ligands and culture dimensionality

To interrogate how air exposure, ECM context and dimensionality interact to shape the behavior of lung tumor epithelium, we established a panel of five culture architectures encompassing 2D and 3D systems under either submerged of ALI conditions. These included conventional monolayer culturing of lung adenocarcinoma (A549) cells on transwell inserts, 2D ALI cultures air-lifted after confluence, 2D_ECM ALI cultures grown on Matrigel-coated inserts under ALI, 3D submerged cultures of A549 cells embedded in Matrigel and 3D ALI culturing maintained with direct apical air exposure (**Figure 1A**). Matrigel was chosen to represent the tumorigenic BM context, as it is enriched in laminin isoforms, collagen IV and heparan sulfate proteoglycans, thereby providing core BM-derived biochemical cues known to regulate epithelial adhesion, polarity and growth [32,33]. Cells were cultured in all models for five weeks, enabling longitudinal evaluation of growth patterns and morphological changes. Brightfield imaging acquired weekly (days 0 to 35) revealed that each culture set-up followed a distinct morphological trajectory driven by its architectural constraints (**Figure 1B**). In 2D submerged conditions, A549 cells expanded as confluent monolayers which gradually thickened into disordered multilayers but did not develop higher-order epithelial patterning. In contrast, 2D ALI and 2D_ECM ALI cultures reorganized noticeably after air-lifting, forming multilayered epithelial structures with conspicuous circular cellular networks which stabilized over time. Similarly, higher magnification images exhibited small air hole structures resembling early ductal patterning and reflecting an ALI-induced reorientation toward airway-like organization. Interestingly, incorporation of ECM cues in the 2D_ECM ALI model further accentuated these changes with more numerous air hole structures compared to 2D ALI alone (**Supplementary Figure 1**). This suggests that BM ligands potentiate ALI-driven epithelial organization through enabling basal anchoring.

**Figure 1.**
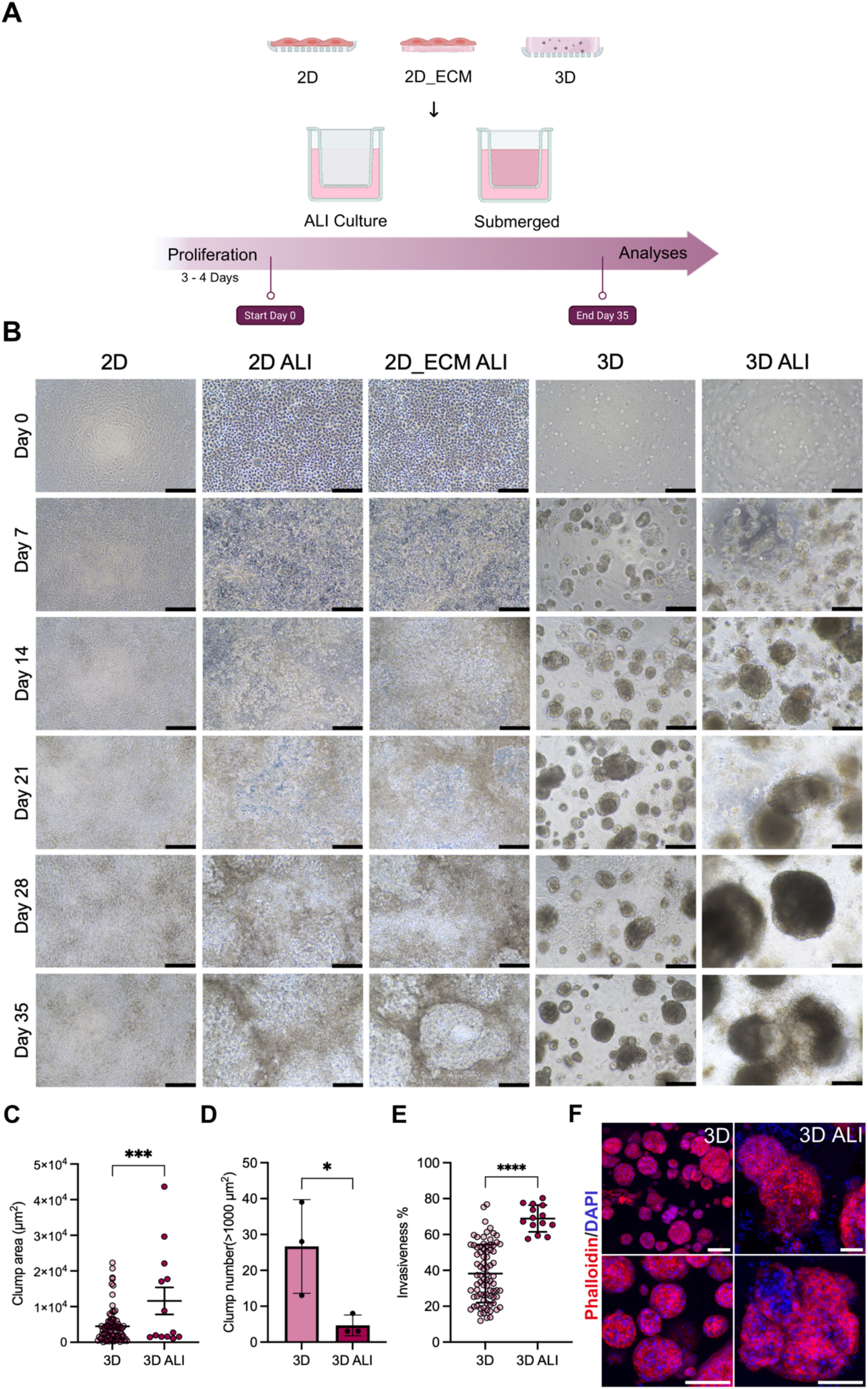
Distinct tumor epithelial states emerge from modulation of air exposure, ECM ligands and culture dimensionality. **A**) Schematic representation of study design (Created via Biorender). **B**) Representative brightfield microscopy images of A549 cells cultured across the five culture set-ups (2D, 2D ALI, 2D_ECM ALI, 3D, 3D ALI) in the study over five weeks. Scale bar: 200 µm. **C**) Quantification of clump area (µm^2^) **D**) clump number (>1000 µm^2^) and **E**) invasiveness (%) of cells grown in different models. Statistical analyses were performed with a two-tailed t-test (n = 3; mean ± s.e.m; ns, no significance, \**p* <0.05, ***p < 0.001, ****p < 0.0001). **F**) Immunofluorescence images of A549 clumps stained for Phalloidin (red) and DAPI (blue) after five weeks of culture in 3D and 3D ALI models. Scale bar: 100 µm.

Dimensionality introduced an additional later of architectural specification. In 3D submerged cultures, A549 cells formed compact spheroids with cohesive organization, morphologies typical for embedded cultures in BM-derived matrices. In striking contrast, 3D ALI cultures yielded substantially larger, structurally heterogenous spheroids exhibiting budding protrusions, irregular borders and outward cellular movement. Air exposure thus exerted a major influence on 3D morphogenesis, promoting both spheroid expansion and invasive remodeling. Quantitative image analyses corroborated these observations. Clump areas observed in 3D ALI conditions were significantly greater than those in 3D submerged cultures (**Figure 1C**). Clump number was significantly higher in the 3D submerged model consistent with many small and contained spheroids rather than the fewer, enlarged forms observed for under ALI (**Figure 1D**). Circularity analysis demonstrated a marked decrease in 3D ALI conditions, indicating greater morphological invasiveness, structural dynamicity and loss of boundary symmetry observed in submerged spheroids (**Figure 1E**). Cytoskeletal organization assessed by phalloidin/DAPI staining further validated these differences (**Figure 1F**). Spheroids in 3D submerged conditions displayed compact structure with distinct contours and densely packed nuclei. In contrast, 3D ALI spheroids exhibited peripheral protrusions and pronounced cellular outgrowth from the spheroidal body, consistent with the invasive morphology detected in brightfield imaging and quantitative shape metrics.

Collectively, these results demonstrate that air exposure, ECM and dimensionality impose distinct constraints on A549 epithelial organization, producing five stable and phenotypically unique microenvironmental states. ALI reorganized 2D cultures into multilayered, airway-like structures, an effect amplified with ECM cues. Dimensionality redefined growth behavior, transforming 3D submerged spheroids into large, invasive and irregular structures when exposed to air. These outcomes establish a foundational basis for understanding the functional, transcriptional and lineage-specific differences in the following sections.

### 2.2 Barrier integrity and permeability as functional outputs across culture models

Having delineated the morphological landscape of the five culture models, we next evaluated their barrier function using TEER measurements and dextran permeability assays. Across the 35-day culture period, 2D submerged monolayers consistently produced the highest TEER values, reaching up to 60 Ω·cm² (**Figure 2A**). These values are lower than what is reported for healthy pulmonary epithelium but align with the low barrier capacity commonly reported for A549 cells in the literature [34]. On the other hand, 2D ALI and 2D_ECM ALI models displayed consistently lower TEER values than 2D submerged cultures at all measured time points. Airway-like morphological reorientation of A549 cells was accompanied by diminished barrier function. Embedded 3D models, whether maintained submerged or under ALI, exhibited uniformly low TEER throughout the experiments, with no significant differences between 3D and 3D ALI. This pattern reflects the limited ability of A549 spheroids to form junction-rich epithelial layers. Overall, TEER analysis showed that barrier function is strongly affected by culture dimensionality, with monolayers producing measurably tighter barriers than 3D aggregates and ALI exposure alone is insufficient to enhance junction formation.

**Figure 2.**
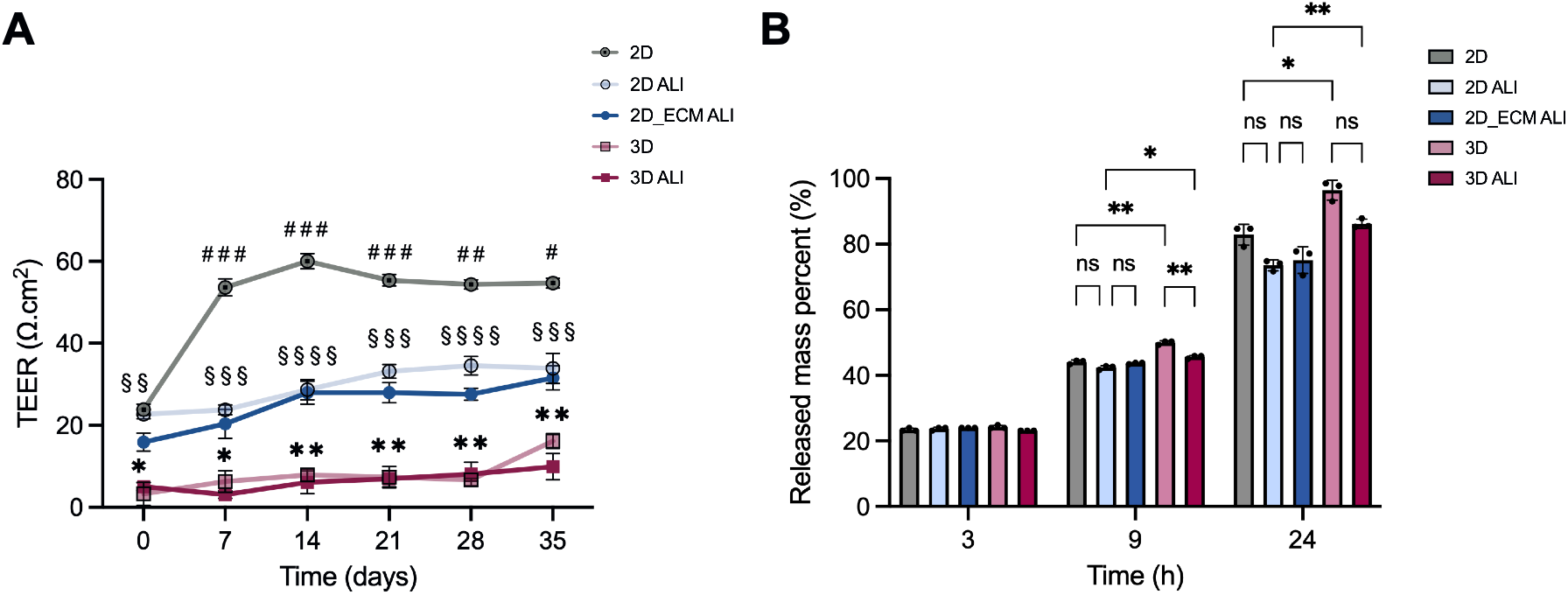
Barrier integrity and permeability as functional outputs across culture models. **A**) Transepithelial electrical resistance (TEER) measurement of cultured samples at day 0, 7, 14, 21, 28 and 35. Data represent the mean ± standard deviation from triplicate samples. *n* = 3; mean ± s.d.; \*\**p* <0.01, \*\*\**p* <0.001 for 2D ALI compared to 3D ALI culture model. ^§§^*p* <0.01, ^§§§^*p* < 0.001, ^§§§§^*p* <0.0001 for 2D compared to 3D culture model. ^#^*p* <0.05, ^##^*p* <0.01, ^###^*p* <0.001 for 2D compared to 2D ALI. **B**) Epithelial barrier permeability at day 35 of culturing measured by Rhodamine B-tagged dextran (10kDa) release from apical to basal chamber of the transwell at 3, 9 and 24 h time points. Data represent mean ± standard deviation from triplicate samples. *n* = 3; mean ± s.d.; ns, no significance, \**p* <0.05, \*\**p* <0.01.

Permeability assessment using 10 kDa Rhodamine B-tagged dextran on day 35 further revealed architectural differences (**Figure 2B**). After 3 hours, all culture conditions released roughly 20% of the tracer with no significant differences, indicating comparable early-phase diffusion. However, dimensionality effects emerged by 9 hours when 3D cultures showed significantly higher dextran release than 2D models. Interestingly, ALI culturing caused a significant decrease in dextran release only in 3D cultures. After 24 hours, differences in permeability among the conditions became more pronounced. 3D conditions demonstrated significantly higher cumulative dextran release than 2D conditions, confirming the weak barrier phenotype of embedded spheroids. On the other hand, ALI culturing significantly attenuated permeability for 3D cultures.

These trends are reproduced in the early time-point data (**Supplementary Figure 2**). At day 7, 3D cultures already exhibit significantly higher permeability than 2D samples after 9 hours. ALI culturing caused reduced release in 3D samples. By 24 hours, permeability differences widen further, with significant increase in dextran release for 3D cultures compared to 2D. Interestingly, although ALI had no impact on release for 2D cultures, ECM incorporation caused a distinct rise at 24 hours.

Taken together, the TEER and dextran assays show that A549 barrier formation is strongly dependent on architectural context. 2D formats produce the tightest and most stable barriers, regardless of ALI exposure or ECM, whereas embedded 3D cultures remain highly permeable with week electrical resistance throughout the 35-day culture period. ALI culturing confers improvements in 3D, though insufficient to approach the barrier performances of 2D conditions. These data confirm that dimensionality, not ALI or ECM alone, acts as the primary determinant of A549 barrier integrity.

### 2.3 Distinct Lineage Signatures Emerge from Modulation of ALI, ECM and Dimensionality

To determine how ALI exposure, ECM context and dimensionality influence epithelial differentiation programs in lung adenocarcinoma, we next quantified a panel of lineage, secretory and junctional markers characteristic of alveolar and airway epithelial identity (**Figure 3**). Among the examined markers, surfactant protein C (SP-C) showed the most striking response across architectures. 3D ALI cultures exhibited a robust and significant upregulation of SP-C expression, whereas all other models remained near baseline. Such selective induction suggests that simultaneous exposure to a 3D ECM microenvironment and apical air is required to re-engage alveolar type II-like transcriptional programs, consistent with reports showing that SP-C expression in transformed lung lines is highly dependent on culture context [35]. Expression of the basal cell-associated transcription factor TP63, a key regulator of airway progenitor activity, revealed a complementary architectural dependence. TP63 expression was significantly higher in ALI conditions compared to submerged ones, with the strongest induction observed for 2D ALI and 2D_ECM ALI cultures. These findings parallel the morphological reorientation seen in **Figure 1**, where ALI drove the emergence of multilayered, air hole-like epithelial structures. Expression of β-tubulin IV (TUBB4A), a hallmark of ciliated cell differentiation, followed a similar profile to TP63 with ALI cultures exhibiting significantly higher expression than submerged ones and with highest levels observed for 2D ALI and 2D_ECM ALI. Although A549 cells are not canonical ciliated progenitors, this upregulation in β-tubulin IV suggests that ALI can partially activate ciliated lineage programs in transformed lung epithelium under both 2D and 3D conditions. On the other hand, the presence of ECM did not exert a significant effect on β-tubulin IV expression. Thyroid transcription factor 1 (TTF-1), a lineage-defining transcription factor required for the maintenance of alveolar identity and retained in adenocarcinomas [36], showed an architecture-sensitive response to ALI exposure. ALI culturing caused a slight upregulation of TTF-1 expression only in the 2D context whereas in 3D embedded models a baseline expression was maintained. Tight junction protein 1 (TJP1/ZO-1) expression remained low across all conditions consistent with the well-established inability of A549 cells to form mature tight junctions. ALI exposure nonetheless induced a modest increase TJP1 expression in both 2D and 3D models. Mucin 5B (MUC5B) revealed a distinct expression profile highly dependent on both ALI and dimensionality. All ALI cultures exhibited significantly higher MUC5B expression compared with their submerged counterparts, consistent with the known role of ALI in promoting mucosecretory differentiation. Notably, 3D ALI model yielded the highest MUC5B expression across all models, exceeding both 2D ALI and 2D_ECM ALI variants despite their pronounced airway-like morphological organization. This indicates that a 3D ECM environment combined with air exposure is a strong driver of mucin induction with high relevance to adenocarcinoma biology where dysregulated mucin production contributes to tumor progression, matrix remodeling and therapeutic resistance [37]. The pronounced MUC5B response in the 3D ALI model parallels the SP-C induction, suggesting that embedded ALI conditions uniquely engage both secretory and alveolar programs not observed in the other models.

**Figure 3.**
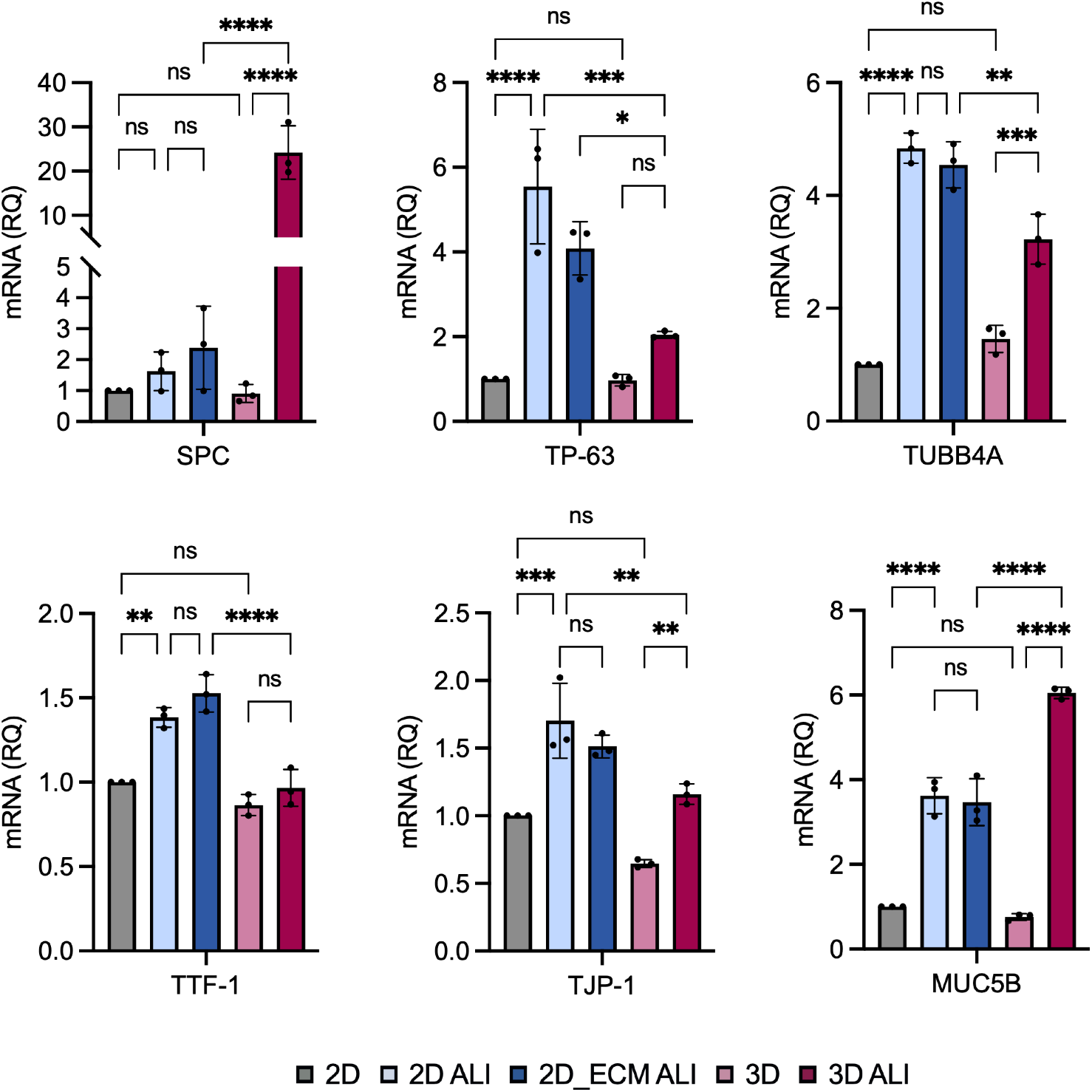
Distinct Lineage Signatures Emerge from Modulation of ALI, ECM and Dimensionality. mRNA expression of lineage markers surfactant protein C (SPC), TP63, β-tubulin IV (TUBB4A), thyroid transcription factor 1 (TTF-1), tight junction protein 1 (TJP1), mucin 5B (MUC5B) in 2D, 2D ALI, 2D_ECM ALI, 3D and 3D ALI conditions. Relative quantitation (RQ) was used with normalization to 2D samples. Data represent mean ± standard deviation from triplicates (ns, no significance, *p < 0.05, **p < 0.01, ***p < 0.001, ****p < 0.0001).

Together, the expression profiles of lineage markers demonstrate that each architectural axis, air exposure, ECM composition and dimensionality, exerts a distinct effect on epithelial differentiation programs in A549 cells. ALI consistently promoted basal and mucociliary lineage markers in line with the airway-like morphologies observed in **Figure 1**. ECM embedding alone did not recapitulated these shifts but potentiated the ALI response when combined in 3D. 3D ALI cultures, providing air exposure, ECM contact and spatial confinement, emerged as the only architecture capable of upregulating both SP-C and MUC5B and reflecting divergent aspects of pulmonary epithelial identity. This dual induction suggests that the interplay between 3D morphology, ECM cues and ALI generates a hybrid state with features of both alveolar type-II and mucosecretory phenotypes which may capture adenocarcinoma plasticity more effectively than traditional 2D or submerged 3D models.

### 2.4 Microenvironment-driven Modulation of Epithelial-Mesenchymal Plasticity

To evaluate whether the architectural cues that shaped morphology and differentiation also altered epithelial-mesenchymal plasticity in A549 cells, we profiled a panel of EMT-associated genes across all five culture configurations (**Figure 4**). Because EMT is tightly linked to invasiveness, progression, lineage plasticity and therapy resistance in lung adenocarcinoma [38], these markers provide functional insight into how ALI, ECM and dimensionality re-pattern tumor cell state. Expression of E-cadherin (CDH1), the canonical epithelial adhesion molecule whose loss denotes the onset of EMT, remained unchanged across all models. This aligns with partial EMT phenotypes reported for lung adenocarcinomas even under strong EMT-stimulating cues and maintenance of E-cadherin while modulating transcriptional EMT regulators [39]. Vimentin (VIM) exhibited modest but significant sensitivity to ALI in 2D conditions. 2D ALI cultures showed an approximately 1.5-fold increase in VIM expression compared to the submerged model, whereas 2D_ECM ALI did not show such increase and ALI exposure even suppressed VIM relative to 2D ALI. This suggests that air exposure alone promotes limited mesenchymal priming in 2D but addition of basement membrane ligands stabilizes epithelial features and reduces ALI-driven VIM induction. Embedding in 3D reduced vimentin expression relative to 2D and ALI did not further alter VIM in 3D, indicating that 3D ECM architecture constrains mesenchymal cytoskeletal remodeling independent of air exposure. ZEB1, a central EMT-inducing transcription factor known to drive polarity loss and invasiveness in lung adenocarcinoma, showed a distinct and more robust architectural response. Both 2D and 3D submerged cultures showed similar baseline expression. However, ALI exposure induced marked increase in ZEB1 expression in 2D cultures, regardless of ECM cues, suggesting that ALI activates mesenchymal programs even when epithelial markers such as E-cadherin remain stable. In 3D, ALI induced a smaller but significant increase in ZEB1 expression compared to 3D submerged cultures. These findings indicate that air exposure is a consistent inducer of ZEB1 with a more pronounced impact in 2D than in 3D supporting the role of apical-basal polarization in amplification of ZEB1-linked plasticity programs [40]. Among all EMT markers profiled, TWIST1 showed the most distinct and architecture-specific response. TWIST1 expression was modestly reduced upon ALI exposure in 2D cultures. Moreover, 3D embedding caused a marked decrease in TWIST1 under submerged conditions. In stark contrast, 3D ALI cultures showed a sharp upregulation of TWIST1, far exceeding all other models. TWIST1 is a potent EMT driver linked to stemness, metastasis and matrix invasion in lung adenocarcinoma [41]. The exclusive and robust activation of TWIST1 expression in 3D ALI suggests that the combination of ECM embedding and apical air exposure engages a strong mesenchymal program, consistent with the invasive spheroid morphologies and reduced circularity quantified in **Figure 1**. N-cadherin (CDH2) remained largely unchanged across 2D and 3D submerged conditions, with only a small but significant decrease observed in 3D ALI relative to 3D submerged and 2D_ECM ALI. This pattern reinforces that A549 cells respond to ALI through fluctuations in transcriptional EMT drivers while N-cadherin and E-cadherin remains largely refractory, consistent with their well-established partial-EMT behavior where transcriptional reprogramming occurs without classical cadherin switching [42–44]. Fibronectin (FN1) expression remained stable across all architectures, indicating that matrix remodeling via fibronectin induction is not a major axis of phenotypic divergence among these conditions. In contrast, snail 1 (SNAI1), another canonical EMT-initiating transcription factor implicated in early polarity loss and invasion, showed strong architectural dependence. In 2D, ALI induced a strong increase in SNAI, further amplified in 2D_ECM ALI, suggesting that both air exposure and basement-membrane cues cooperate to activate SNA1-dependent epithelial plasticity. In contrast, 2D and 3D submerged cultures showed similarly low SNA1 expression, demonstrating that dimensionality alone does not alter this program. Consistent with the TWIST1 surge, 3D ALI also produced a sharp increase in SNA1 relative to 3D submerged. Finally, α-smooth muscle actin (α-SMA/ACTA2) demonstrated a highly architecture-specific pattern. 2D ALI strongly upregulated ACTA2 expression relative to 2D submerged, however, 2D_ECM ALI potently reduced ACTA2 expression back to 2D baseline, indicating that basement-membrane cues suppress ALI-induced myofibroblast-like activation. In 3D, ACTA2 remained at basal levels regardless of ALI exposure, suggesting that embedded architecture constrains ACTA2 induction even when TWIST1 and SNAI1 are strongly activated.

**Figure 4.**
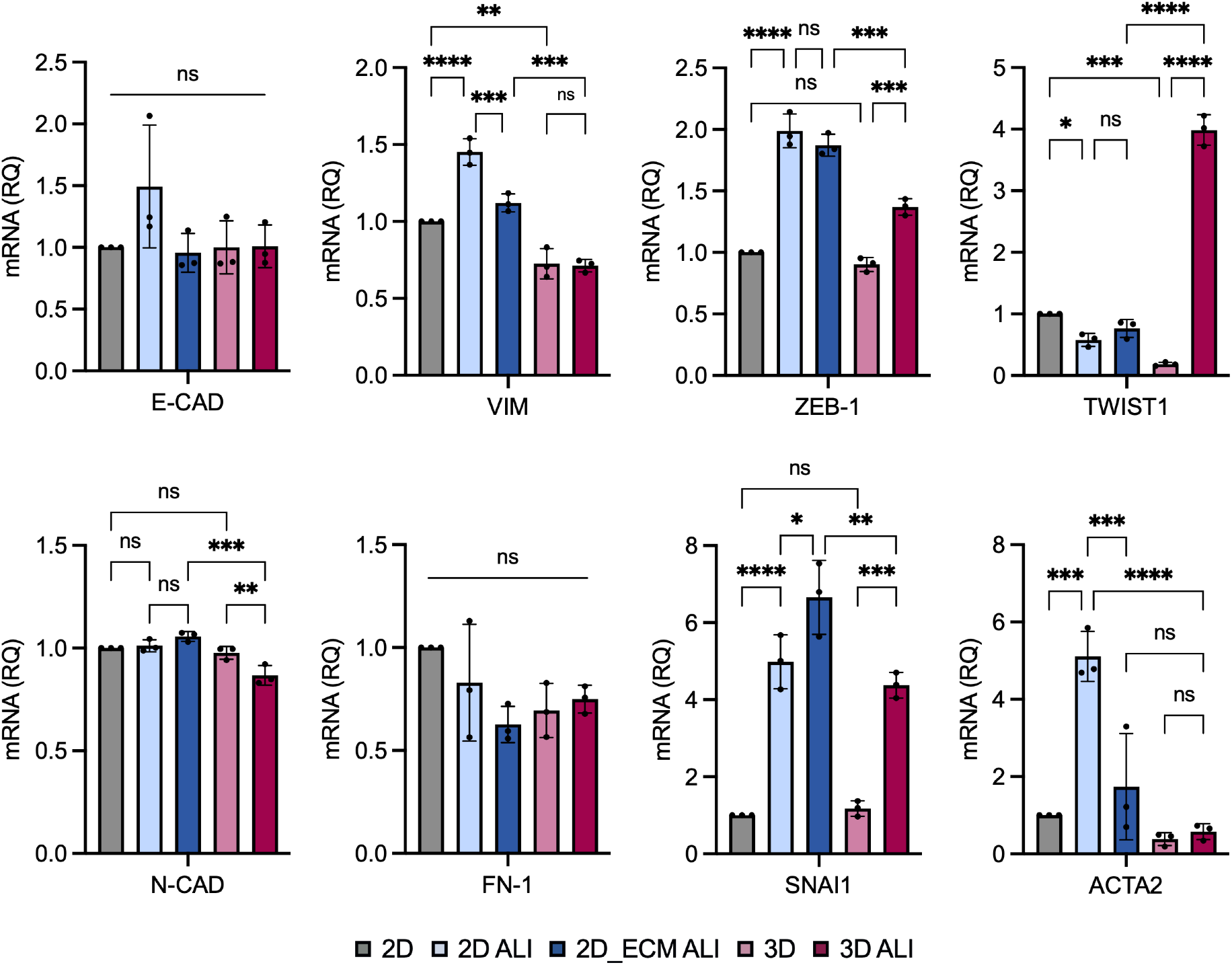
Microenvironment-driven Modulation of EMT Transcriptional Plasticity. mRNA expression of E-cadherin (E-CAD), vimentin (VIM), ZEB-1, TWIST1, N-cadherin (N-CAD), fibronectin (FN-1), snail 1 (SNAI1) and α-smooth muscle actin (α-SMA/ACTA2) in 2D, 2D ALI, 2D_ECM ALI, 3D and 3D ALI conditions. Relative quantitation (RQ) was used with normalization to 2D samples. Data represent mean ± standard deviation from triplicates (ns, no significance, *p < 0.05, **p < 0.01, ***p < 0.001, ****p < 0.0001).

Collectively, these data show that ALI is a dominant inducer of EMT-linked transcriptional reprogramming in lung adenocarcinoma while dimensionality modulates the depth and nature of the response. 2D ALI primarily induces partial EMT with strong ZEB1, SNAI1 and ACTA2 activation, whereas 3D ALI uniquely activates TWIST1-driven invasive programs consistent with the aggressive spheroid morphologies. ECM cues in 2D selectively suppresses ALI-driven VIM and ACTA2 expression, while ECM embedding in 3D maintains low junctional strength yet potentiates TWIST1 and mucin induction under ALI (**Figures 2-4**). These layered transcriptional outcomes illustrate that ALI, ECM and dimensionality converge to generate distinct, non-linear EMT states reflecting the hybrid and context-dependent EMT phenotypes characteristic of lung adenocarcinoma [38].

### 2.5 Architectural modulation of inflammatory signatures

To assess whether microenvironmental architecture modulates inflammatory signaling and matrix remodeling programs in lung adenocarcinoma, we analyzed expression of interleukin-6 (IL-6), interleukin-8 (IL-8) and matrix metalloproteinase-12 (MMP-12), cytokines widely expressed in NSCLC and associated with tissue-level inflammation, tumor proliferation and progression [45], across all five culture configurations (**Figure 5**). IL-6 demonstrated pronounced sensitivity to both ALI exposure and dimensionality. In 2D cultures, ALI induced a marked upregulation relative to 2D submerged cultures indicating that apical air exposure elevates pro-inflammatory signaling. Basal ECM incorporation on 2D under ALI conditions did not exert an effect. On the other hand, 3D ECM embedding under submerged conditions caused a distinct increase in IL-6 expression compared to 2D. 3D ALI culturing triggered a sharp increase relative to both 3D submerged and 2D ALI conditions. IL-8, a pro-inflammatory cytokine which correlates with poor prognosis, immune evasion and aggressive phenotypes [46,47], similarly showed a strongly ALI-induced expression in both 2D and 3D settings. However, 2D ALI and 3D ALI did not differ significantly, suggesting that IL-8 is less sensitive to architectural cues compared to IL-6. Collectively, IL-6 and IL-8 expression profiles show that air exposure is a robust enhancer of pro-inflammatory cytokine expression across both dimensional contexts. MMP-12, a metalloproteinase linked to lung remodeling and tumor progression [48,49], directly activates IL-8 secretion in lung adenocarcinoma and participates in pulmonary inflammatory programs [50]. In contrast with IL-6 and IL-8 profiles, MMP-12 expression showed a distinct architectural dependency in its response to ALI. ALI strongly induced MMP-12 expression only in 3D cultures revealing a specific synergy between ECM embedding and air exposure that is not recapitulated in 2D.

**Figure 5.**
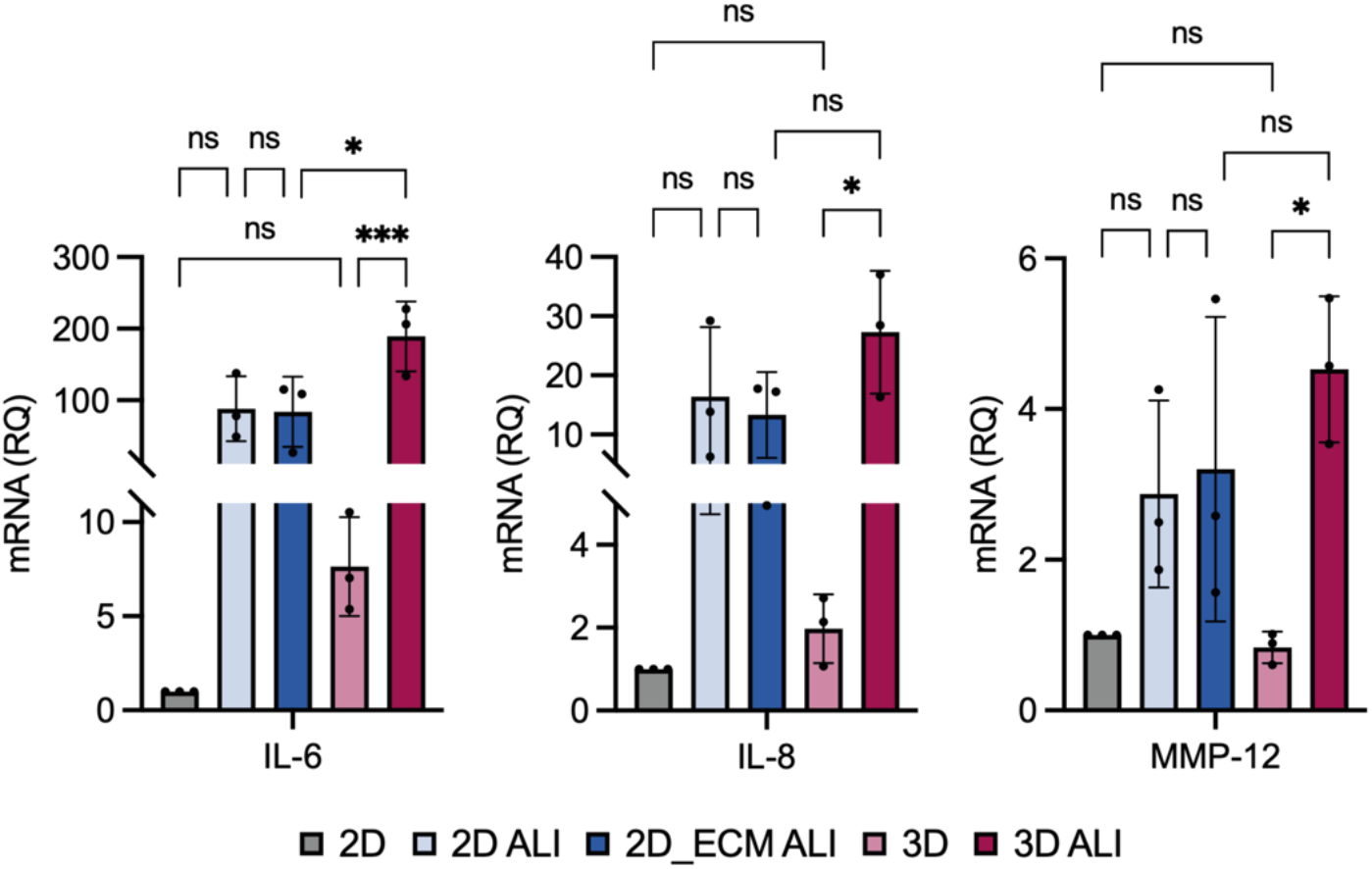
Architectural modulation of inflammatory signatures. mRNA expression of interleukin-6 (IL-6), interleukin-8 (IL-8) and matrix metalloproteinase-12 (MMP-12) in 2D, 2D ALI, 2D_ECM ALI, 3D and 3D ALI conditions. Relative quantitation (RQ) was used with normalization to 2D samples. Data represent mean ± standard deviation from triplicates (ns, no significance, *p < 0.05, **p < 0.01, ***p < 0.001, ****p < 0.0001).

Taken together, the inflammatory and remodeling markers follow a coherent architectural pattern which shows that ALI is the dominant inducer of IL-6 and IL-8 expression, triggering a strong inflammatory response in both 2D and 3D contexts. 3D embedding further amplifies ALI-driven response and shows that matrix-anchored cultures with apical air generate more complex inflammatory and remodeling signatures.

### 2.6 Architectural regulation of epithelial identity, progenitor state and mucosecretory function

To visualize how microenvironmental architecture shapes epithelial organization and lineage allocation at protein level, we performed immunofluorescence stainings for cytoskeletal structure (phalloidin), epithelial identity (EpCAM), basal/progenitor status (p63) as well as mucosecretory (MUC5B) and ciliated (β-tubulin IV) function across all five culture models (**Figures 6-7**). Consistent with the morphological trajectories observed in brightfield imaging (**Figure 1**), phalloidin staining demonstrated that 2D submerged cultures formed confluent, planar layers while air exposure led to enhanced reorganization and epithelial folding. In 3D models, spheroids displayed dense cell aggregates under submerged conditions whereas 3D ALI spheroids exhibited irregular, protrusive extensions that mirror their invasive morphology. Staining for EpCAM revealed that 2D submerged cellular layers expressed largely fail to adopt epithelial identity except for a few localized regions, consistent with their heterogenous organization. In contrast, 2D ALI and 2D_ECM ALI cultures displayed uniform EpCAM expression, reflecting strengthened epithelial polarity and cohesiveness under air exposure. 3D submerged spheroids did not express EpCAM, where 3D ALI conditions produced strongly EpCAM-positive clusters throughout, illustrating that apical air exposure in 3D fosters a robust epithelial identity of tumor cells otherwise absent in submerged matrices. Basal/progenitor activity, assessed with p63 expression, also demonstrated strong architectural dependence. 2D submerged cultures showed negligible p63 expression whereas ALI exposure led to widespread and intense p63 expression. However, in 2D_ECM ALI, p63 was still induced but became spatially restricted to discrete cellular clusters rather than diffusely expressed, suggesting that basement-membrane ligands modulate how ALI-driven basal programs are deployed. In 3D submerged cultures, p63 expression was largely absent. Strikingly, 3D ALI cultures showed abundant p63-positive cells distributed throughout the invasive epithelial clusters seen in this model demonstrating that apical air exposure in a matrix-embedded context reawakens progenitor programs. Together, the stainings support the gene expression data in Figure 3 and emphasize that ALI is a key driver of basal lineage activation across dimensional contexts.

**Figure 6.**
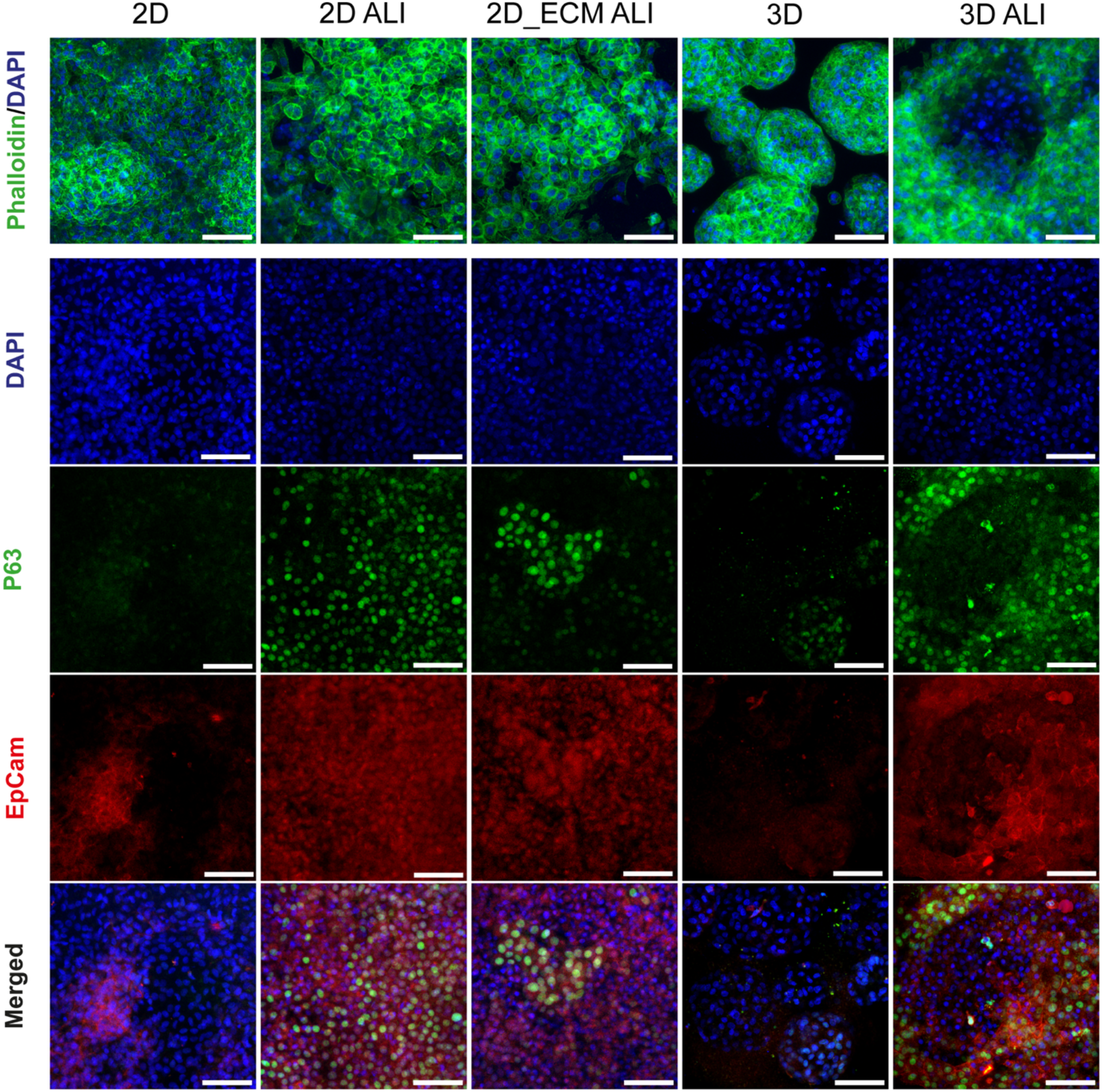
Architectural regulation of epithelial identity and progenitor state. Immunofluorescence stainings of 2D, 2D ALI, 2D_ECM ALI, 3D and 3D ALI cultures. The upper panel shows actin organization visualized by phalloidin and nuclear staining with DAPI. The lower panel displays separate and merged channels for p63, EpCAM and DAPI stainings. Scale bars: 100 μm.

**Figure 7.**
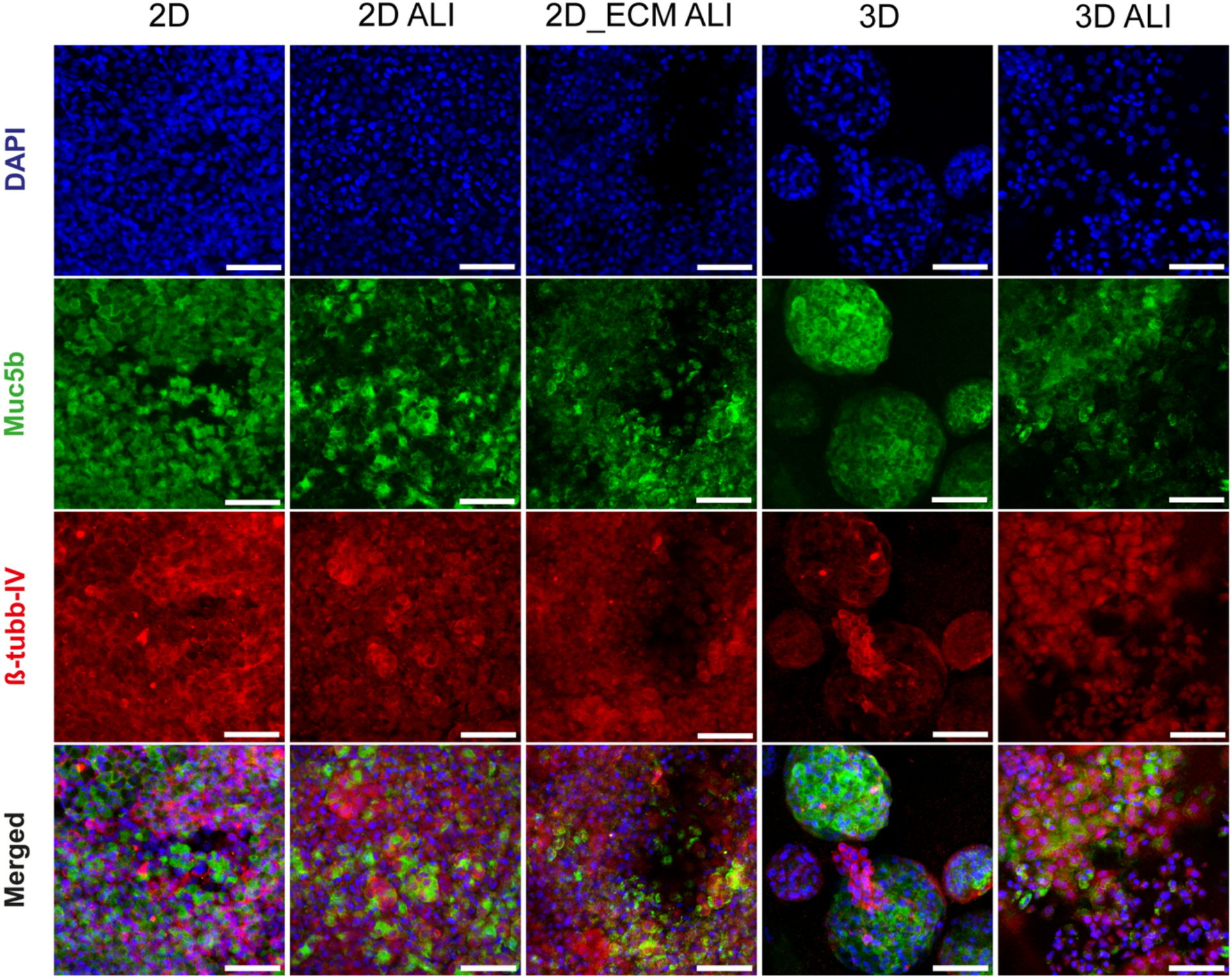
Architectural regulation of mucosecretory function. Immunofluorescence staining images of 2D, 2D ALI, 2D_ECM ALI, 3D and 3D ALI cultures for MUC5B and β-tubulin IV. DAPI was used for nuclei counterstaining. Scale bars: 100 μm.

MUC5B and β-tubulin IV stainings (Figure 7) revealed further architectural specialization. MUC5B expression was broadly distributed across cells in 2D submerged monolayers, whereas 2D ALI cultures displayed fewer but sharply defined MUC5B-positive cells. 2D_ECM ALI cultures showed a more clustered MUC5B expression with groups of neighboring cells stained strongly, contrasting with the more dispersed expression pattern in 2D ALI and indicating that ECM ligands support the emergence of localized secretory foci under ALI. 3D submerged cultures showed heterogenous MUC5B expression with some spheroids uniformly positive and others negative. Notably, in several MUC5B-positive spheroids, peripheral expression of β-tubulin IV was observed. Cellular extensions emanating from MUC5B-positive spheroids were strongly β-tubulin IV-positive but MUC5B-negative, suggesting functional segregation between spheroid core and migratory fronts. In 3D ALI cultures, large and invasive spheroids displayed co-localization of markers rather than segregation where uniform β-tubulin IV staining and large areas of integrated MUC5B staining was observed. These findings align with the transcriptional enhancements of MUC5B and β-tubulin IV observed in 3D ALI (**Figure 3**). Collectively, the immunofluorescence data demonstrate that ALI is a central cue driving basal activation, epithelial identity and mucosecretory function whereas ECM embedding in 3D modulates how these programs are spatially organized.

## 3. Discussion

Lung adenocarcinoma exists not as a fixed epithelial entity but as a collection of plastic, context-dependent states which emerge through interactions with microenvironmental cues. Conventional in vitro systems such as submerged monolayers or 3D spheroids capture limited aspects of this landscape due to modulating architectural features along a single axis, masking emergent states which arise upon their intersection. By systematically varying air exposure, ECM ligands and dimensionality, this study shows that these axes constitute a powerful architectural triad and act in combinatorial and nonlinear ways to unlock epithelial, progenitor, inflammatory and mesenchymal programs in A549 cells. Although A549 is often portrayed as a static ATII-like model, our findings reveal that its phenotype is highly contingent on the structural and biochemical environment uniquely eliciting hybrid and clinically relevant behavior. The resulting architectural phenotypes provide new insights into how lung adenocarcinoma interprets physical context and highlight design principles for more physiologically anchored tumor models.

A central insight in the study is the dominant instructive role of ALI in epithelial organization, progenitor activation, mucosecretory programs, transcriptional EMT drivers and inflammatory markers across the models. In contrast to submerged cultures, ALI reorganized tumor cells into cohesive epithelial structures with strongly induced EpCAM expression indicating recovery of epithelial identity typically eroded in conventional dysplastic monolayers. The robust epithelial marker expression alongside with an invasive morphology and activation of EMT markers under ALI conditions supports the emerging view that adenocarcinomas occupy hybrid epithelial states which are neither fully polarized nor fully mesenchymal and that the microenvironment can shift the balance of such plasticity [38]. In parallel, ALI exposure caused potent activation of p63 expression. Basal-like transcriptional programs are increasingly associated with aggressive disease, poor clinical outcome and therapeutic resistance in NSCLC [51]. Our findings that p63 is virtually absent in submerged models but strongly induced in both 2D and 3D ALI contexts suggest that air exposure enables a niche that lung tumors can exploit to activate progenitor-like states. This interpretation is reinforced by reports linking progenitor-like and lineage-plastic epithelial states to lung adenocarcinoma progression, while epithelial plasticity has also been implicated in therapeutic resilience [52–54]. The spatial restriction of p63-positive cells into discrete clusters in the ECM-anchored 2D ALI model as opposed to widespread expression of p63 in 2D ALI implies that basement-membrane ligands alter patterning of ALI-driven progenitor response. Such patterning of p63 is consistent with tumor histology where basal-like foci can arise within otherwise alveolar tumors [51, 52]. In contrast, 3D ALI generated global p63 expression across invasive spheroids indicating that matrix embedding synergizes with ALI to create a niche permissive to progenitor-dominated, regenerative-like tumor states.

Equally compelling is the convergence of alveolar and mucinous signatures in 3D ALI where SP-C and MUC5B were co-induced upon air exposure, phenotypes seldomly occur together in standard cultures. SP-C expression is typically diminished in transformed ATII-derived lines in the absence of differentiation cues [35], thus its re-activation in 3D ALI suggests recapitulation of the microenvironmental elements of a regenerative alveolar niche. Conversely, MUC5B is linked to mucinous adenocarcinoma, KRAS-driven tumor progression and matrix remodeling [27, 37]. The coexistence of alveolar and mucinous signatures, normally segregated along distinct epithelial lineages, suggests a hybrid state increasingly recognized in aggressive LUAD subtypes with high plasticity and metastatic potential, exhibiting both secretory expansion and self-renewal [52–54]. This state is not recaptured in 2D ALI, 2D_ECM ALI or submerged cultures, underscoring that synergy of matrix embedding and ALI is required to unlock this hybrid differentiation. β-tubulin IV further illustrates the architectural dependence of tumor cell identity. Although conventionally linked to ciliated epithelium, TUBB4A was shown to act on the EMT-cytoskeleton regulatory axis and contribute to motility and metastatic competence in cancer-context [55]. In submerged 3D cultures, β-tubulin IV concentrates peripherally around MUC5B-positive spheroids and at the migratory fronts. However, in 3D ALI, β-tubulin IV expression becomes dispersed throughout invasive clusters indicating a shift toward global cytoskeletal reorganization. This aligns with 3D ALI-specific TWIST1-high transcriptional states, which have been linked to microtubule rearrangement, anchorage-independent migration and metastasis [40,41,43]. The distinct upregulation of TWIST1 suggests a deeply invasive mesenchymal program and is reinforced with the ALI-induced upregulation of other EMT-associated transcriptional drivers such as ZEB1 and SNA1. Yet, classical EMT endpoints, loss of E-cadherin and gain of N-cadherin, remained unchanged. This confirms that A549 cells occupy partial EMT states, as observed in many patient-derived adenocarcinomas, where transcriptional reprogramming rather than cadherin switching defines the EMT spectrum [43,44].

Inflammatory signaling further supports an ALI-responsive, tumor-promoting phenotype. IL-6 and IL-8, key cytokines associated with tumor progression, angiogenesis and metastatic niche formation [45–47], were strongly induced by ALI across both dimensionalities, indicating a shift of tumor cells toward a pro-inflammatory state upon air exposure. MMP-12, a matrix remodeling enzyme linked to metastatic dissemination [48,49], responded exclusively in 3D ALI which highlights that ECM engagement with air exposure is required to unlock matrix-degradative and stromal-interactive states. These inflammatory signatures align with the hybrid epithelial-mesenchymal-secretory phenotypes observed and may reflect a coordinated architectural programming of tumor aggressiveness.

Together, these findings elucidate that air exposure, ECM and dimensionality act as combinatorial regulators that enable distinct tumor states of which 2D or submerged 3D cultures capture a limited subset. In contrast, 3D ALI cultures, combining air exposure, matrix ligands and spatial confinement, reproduces multiple hallmarks of aggressive adenocarcinoma including invasive morphology, progenitor activation, mucin overproduction, cytoskeletal reorganization, hybrid EMT states and heightened inflammatory and remodeling responses. While this work provides architectural insights for lung tumor models, several limitations offer opportunities for future investigation. As A549 cells capture a subset of LUAD diversity, implementing patient-derived organoids would allow studying how the response to architectural cues vary with genotypic diversity. Integrating co-cultures with stromal and immune cells would enable modeling cellular interactions that strongly modulate inflammatory signaling and EMT plasticity in vivo [20, 21]. Incorporation of engineered biomaterials with controlled tunability of mechanical properties and biochemical ligands would permit further dissection of how ECM characteristics synergize with ALI. Nevertheless, the work establishes a framework for microenvironmentally grounded in vitro respiratory tumor models with ability to reshape tumor epithelial identity and plasticity. As next-generation organoids, lung-on-chip platforms and precision oncology approaches increasingly demand physiological relevance, these findings identify ALI-ECM-dimension intersections as critical design parameters for studying tumor behavior.

## 4. Materials and Methods

### 4.1 Cell culture

Human lung adenocarcinoma cell line, A549 (CCL185), was purchased from American Tissue Culture Collection (ATCC). Cells were cultured in DMEM/F12 (Lonza) growth medium supplemented with 10% fetal bovine serum (FBS) (Biowest) and 1% penicillin-streptomycin (P/S) (Biowest) with incubation at 37 °C and 5% CO_2_. Prior to seeding across the culture models in the study, cells were propagated, trypsinized, washed and resuspended. For all models, 24-well transwell inserts with 0.4 pore diameter (Sarstedt, Cat# 83.3932.041) were used for culturing. For 2D, 2D ALI and 2D_ECM ALI set-ups, A549 cells were seeded on the transwell membranes at a density of 2×10^4^ cells per well. In 2D_ECM ALI model, the transwell inserts were coated with 1:20 diluted Matrigel (Corning, Cat# 354230) and incubated for an hour prior to cell seeding. For the 3D models, cells were encapsulated in Matrigel at a final density of 2×10^4^ cells per gel which was casted directly on the transwell membranes. Transwell cultures were incubated at 37 °C for gelation and growth medium was added afterwards. Before the start of air-liquid interface culture, all models were cultured under submerged conditions for a 3-4 day propagation period in which the apical and basal chambers were filled with growth medium.

### 4.2 Air-liquid interface (ALI) culture

Upon reaching 80-90% confluency during propagation, the ALI models (2D ALI, 2D_ECM ALI, 3D ALI) were air-lifted through gentle removal of the culture medium from the apical chambers of the transwell membrane inserts. Submerged cultures (2D, 3D) were kept in growth medium in both chambers. Growth medium was replaced every 3 days. For ALI cultures, the media in the basal chambers were replaced while the apical chambers were washed with D-PBS (Biowest, Cat# L0615-500). All models were cultured up to 5 weeks in humidified incubators at 37 °C and 5% CO_2_ after which samples were collected for downstream analyses.

### 4.3 Transepithelial electrical resistance (TEER) measurement

To measure the integrity of epithelial barriers and junction formation in different culture models, transepithelial electrical resistance (TEER) measurements were performed. Measurements were done weekly during the 5-week culture period on all models in the study (2D, 2D ALI, 2D_ECM ALI, 3D, 3D ALI). EVOM2 ohm meter (Millicell ERS-2, EVOM, Merck Millipore) was used to assess TEER in different models. The resistance of empty transwell membrane was recorded as the blank value (R_blank_) and the resistance of cellular layers on transwell membranes in all conditions were measured as total resistance (R_total_) in units of Ω. The resistance of cell layer was calculated as [54]:

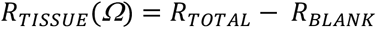

The measurement of resistance is inversely proportional to the area of transwell membrane, so the final reported TEER value was calculated as:

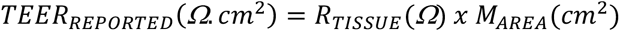

### 4.4 Barrier permeability assay

Barrier permeability characterization was performed by measuring the rate of rhodamine B-tagged dextran (10 kDa) (Invitrogen, Cat# D1824) transport from the apical chamber to the basal chamber of transwell membranes. At designated time points (day 7 and day 35), the samples were collected, the culture medium was removed from both chambers and rinsed gently with D-PBS (Biowest, Cat# L0615-500) at room temperature. Apical chamber was filled with a 0.1 mg/mL of dextran solution in D-PBS and the basal chamber was filled with plain D-PBS. Sampling was done from the basal chamber after 3, 9 and 24 hours. Dextran content was quantified with fluorescence readout using a microplate reader at 570 nm/590 nm excitation/emission. Cumulative release was calculated and reported. At least three replicates were used for each condition.

### 4.5 RT-qPCR analysis

Total RNA from samples was extracted through NucleoSpin^®^ RNA isolation kit (Macherey-Nagel) according to the manufacturer’s protocol. Complementary DNA was synthesized using equivalent amounts of total RNA by using M-MLV Reverse Transcriptase kit (Thermo Fisher, Cat#28025013) according to the manufacturer’s instructions. qRT-PCR was performed using Sybr Green I Master (Roche Diagnostics, Cat#04887352001) with LightCycler^®^ 480-II Instrument (Roche Diagnostics). The sequences of primers are given in **Supplementary Table 1**. All experiments were performed in triplicates and relative quantitation of gene expression was performed with the 2^-ΔΔCt^ method where GAPDH was used as the housekeeping gene.

### 4.6 Immunofluorescence Staining

All transwell cultures were washed with D-PBS and fixed with 4% paraformaldehyde at room temperature for 20 min and stored at 4°C in D-PBS until staining. Cells were permeabilized with 0.1% Triton X-100 in D-PBS and blocked with 5% normal goat serum (Thermo Fisher, Cat#16210064) and 0.1% Triton X-100 in D-PBS for 1 h at room temperature. Then, cells were incubated with primary antibodies in 1% BSA (Sigma Aldrich), 1% normal goat serum and 0.1% Triton X-100 in D-PBS overnight at 4°C. Primary antibodies: EpCam (Biolegend, Cat#324201; 1:200), p63 (CST, Cat#13109; 1:200), Muc5B (Sigma, Cat#HPA008246; 1:100) and β-tubulin IV (Sigma, Cat#T7941; 1:100). After primary antibody incubation, the samples were washed three times with 0.1% Triton X-100 in D-PBS and incubated with diluted secondary antibodies in 1% BSA, 1% normal goat serum and 0.1% Triton X-100 in D-PBS for 1 hour at room temperature. Secondary antibodies: Alexa Fluor 488 goat anti-mouse (Invitrogen, Cat#A11029; 1:1000) and Alexa Fluor 594 goat anti-rabbit (Invitrogen, Cat#R37117; 1:1000). After secondary antibody incubation, the samples were washed again three times with 0.1% Triton X-100 in D-PBS and counterstained for 5 min with DAPI (1 µg/ml in D-PBS) (Sigma). The samples were washed 4 times for 10 min with D-PBS and mounted in Vectashield mounting media (Vector Laboratories). Visualization was performed with confocal microscopy (Leica DMI8-SP8).

### 4.7 Morphological assessments

To monitor the morphological changes of A549 cells in different set-ups, samples were collected at day 35 and fixed with paraformaldehyde (4%) for 20 min at room temperature. Cells were permeabilized with 0.1% Triton X-100 in D-PBS and blocked with 5% normal goat serum (Thermo Fisher, Cat#16210064) and 0.1% Triton X-100 in D-PBS for 1 h at room temperature. F-actin staining was performed using Phalloidin-iFluor Reagent (1:1000, Abcam) for 45 min at room temperature after which samples were thoroughly washed. Nuclei staining was performed with DAPI (1 µg/mL, Sigma) and samples were washed twice for 10 min before imaging. Samples were imaged with Leica SP8 confocal microscope in z-stacks with a 5 μm step length for at least 3 hydrogels. Clump area, clump number and invasiveness analyses were performed as previously reported [56] with ImageJ 2.14.0. Invasiveness of clumps were assessed with the circularity output from the cluster size analysis where a circularity value of 1 indicates a perfectly circular cellular clump with 0% invasiveness whereas a value of 0 circularity indicates 100% invasiveness.

### 4.8 Statistical analysis

Data were analyzed using GraphPad Prism 10.3.0. Quantitative data were presented as mean ± standard deviation. Statistical analyses were performed with one-way ANOVA with Tukey test for pairwise comparisons on multiple groups. A *p*-value of <0.05 was regarded as statistically significant.

## Supporting Information

Supporting Information is available.

## Acknowledgements

The authors acknowledge funding from the International Fellowship for Outstanding Researchers Program of TÜBİTAK (118C238). The entire responsibility of the publication belongs to the owner, the financial support from TÜBİTAK does not mean that the content of the publication is approved in a scientific sense by TÜBİTAK. BioRender.com was used to create the scheme in Figure 1. The authors gratefully acknowledge the use of services and facilities of Koç University Research Center for Translational Medicine (KUTTAM).

## Author Contributions

E.Ö. conceived the project, obtained funding and designed the research. K.Y. conducted experiments and analyzed data. K.Y. drafted and E.Ö. wrote the manuscript.

## Data Availability Statement

The data supporting the findings of the study are provided in the manuscript, its Supplementary Information and from the corresponding author upon reasonable request.

