## Supplementary Information for "Matrix-modulated Air-liquid Interface Microenvironments Reveal Architectural Control of Lung Tumor Phenotypes"

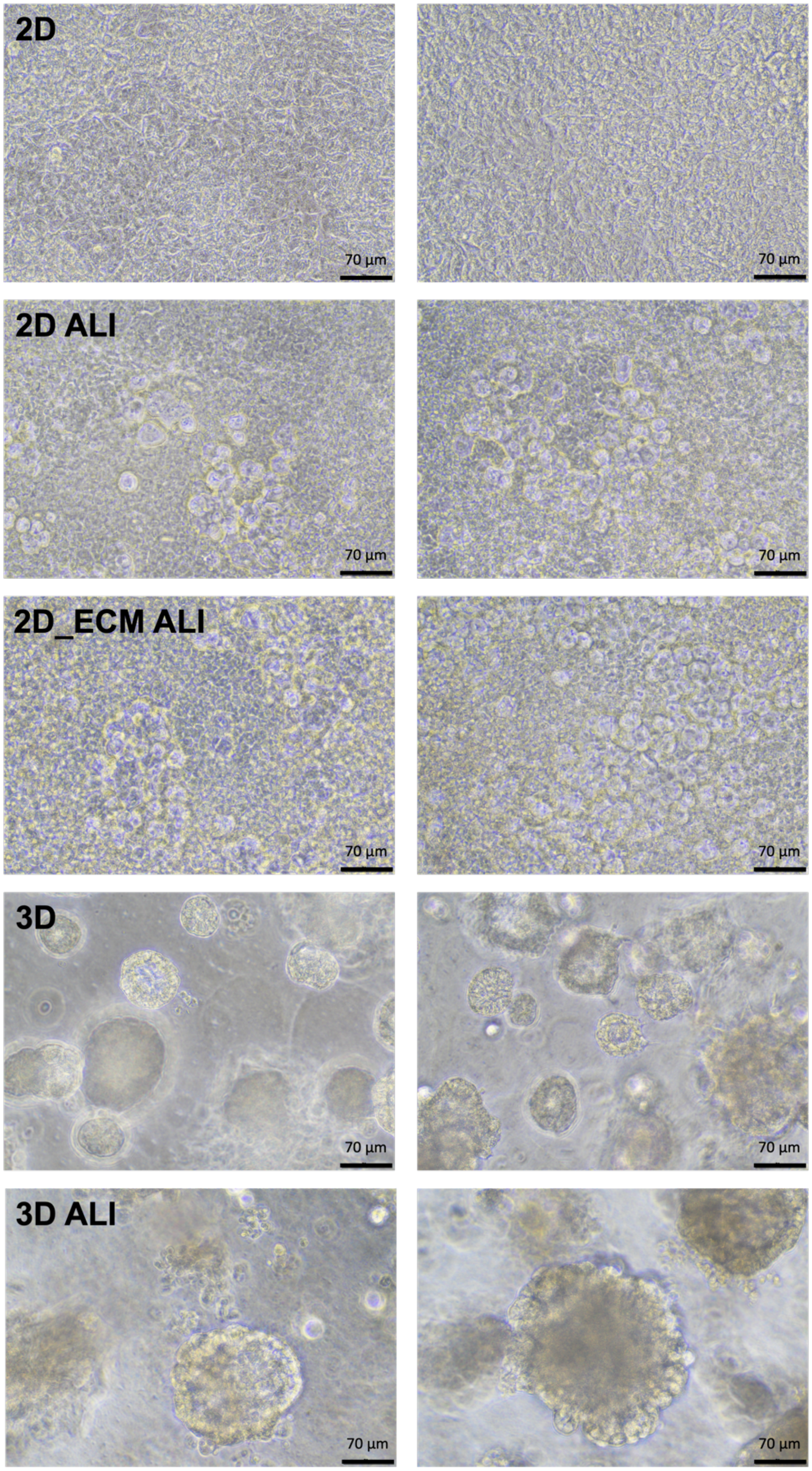


**Supplementary Figure 1.** Representative brightfield microscopy images of A549 cells cultured across the five culture set-ups (2D, 2D ALI, 2D_ECM ALI, 3D, 3D ALI) in the study. Scale bar: 70 µm.

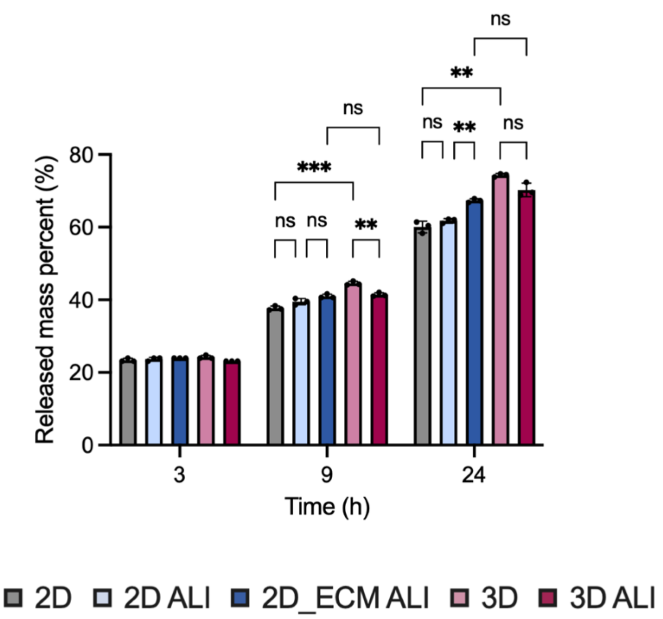


**Supplementary Figure 2.** Epithelial barrier permeability at day 7 of culturing measured by Rhodamine B-tagged dextran (10kDa) release from apical to basal chamber of the transwell at 3, 9 and 24 h time points. Data represent mean ± standard deviation from triplicate samples. $n$ = 3; mean ± s.d.; ns, no significance, $^{**}p$ <0.01, $^{***}p$ <0.001.

**Table 1** The sequences of primers used for qRT-PCR.

| Gene | Forward Primer | Reverse Primer |
|---|---|---|
| SP-C | 5'-GGTGCCGCTCTACTACATCT-3' | 5'CTCCCCAAGCTCCCATTTCT-3' |
| TTF-1 | 5'-TTGGTAGGAACATTGCCCG-3' | 5'-CGTCTTCCAGTCATTCCCAAG-3' |
| TJP-1 | 5'-GATCTGCATCCTTAGAGACCAAG-3' | 5'-CTGAGAAGTGGGTTTGGGAC-3' |
| MUC5B | 5'-CTACATCAAGGTCAGCATCCG-3' | 5'-ATAGAACTCGTTGAAGGCCG-3' |
| TUBB4A | 5'-CACCCAGCAGATGTTCGATG-3' | 5'-CTTCACGTTGTTGGGGATCC-3' |
| TP-63 | 5'-GCCATGCCCAGTATGTAGAAG-3' | 5'-TCATCCCTCCAACACAACTG-3' |
| IL-6 | 5'-CAACCTGAACCTTCCAAAGATG-3' | 5'-ACCTCAAACTCCAAAAGACCAG-3' |
| IL-8 | 5'-AGCCTTCCTGATTTCTGCAG-3' | 5'-GTCCACTCTCAATCACTCTCAG-3' |
| MMP-12 | 5'-CAACTGGACACATCTACCCTG-3' | 5'-ACGGTTCATGTCAGGTGTG-3' |
| E-CAD | 5'-CTGCCAATCCCGATGAAATTG-3' | 5'-TCCTTCATAGTCAAACACGAGC-3' |
| N-CAD | 5'-CAGAATCAGTGGCGGAGATC-3' | 5'-CAGCAACAGTAAGGACAAACATC-3' |
| VIM | 5'-ACCAGCTAACCAACGACAAAG-3' | 5'-AAAGATTGCAGGGTGTTTTCG-3' |
| FN-1 | 5'-CAGAGGCATAAGGTTCGGG-3' | 5'-TTCAGACATTCGTTCCCACTC-3' |
| ZEB-1 | 5'-ACCCTTGAAAGTGATCCAGC-3' | 5'-CATTCCATTTTCTGTCTTCCGC-3' |
| SNAIL | 5'-GGAAGCCTAACTACAGCGAG-3' | 5'-CAGAGTCCCAGATGAGCATTG-3' |
| SLUG | 5'-AGCATTTCAACGCCTCCA-3' | 5'-GGATCTCTGGTTGTGGTATGAC-3' |
| TWIST-1 | 5'-GCATTCTCAAGAGGTCGTGC-3' | 5'-TGACTATGGTTTTGCAGGCC-3' |
| $\alpha$-SMA | 5'-AGCCCTCCTTCATCGGTATG-3' | 5'-TGATCTTCATGGTGCTGGGT-3' |
| GAPDH | 5'-CTGACTTCAACAGCGACACC-3' | 5'-GTGGTCCAGGGGTCTTACTC-3' |
